# Deciphering mixed infections by plant RNA virus and reconstructing complete genomes simultaneously present within-host

**DOI:** 10.1101/2024.02.29.582683

**Authors:** Martine Bangratz, Aurore Comte, Estelle Billard, Abdoul Kader Guigma, Guillaume Gandolfi, Abalo Itolou Kassankogno, Drissa Sérémé, Nils Poulicard, Charlotte Tollenaere

## Abstract

The local co-circulation of multiple phylogenetic lineages is particularly likely for rapidly evolving pathogens in the current globalization context. When various phylogenetic lineages co-occur in the same fields, they may simultaneously be present in the same host plant (i.e. mixed infection), with potential important consequences for disease outcome. This is the case in Burkina Faso for the rice yellow mottle virus (RYMV), endemic to Africa, where it constitutes a major constraint to rice production.

We aimed at deciphering the distinct RYMV isolates simultaneously infecting a single rice plant and sequencing their genomes. To this purpose, we tested various sequencing strategies, and we finally combined direct cDNA ONT (Oxford Nanopore Technology) sequencing with the bioinformatics tool RVhaplo. This methodology was validated though the successful reconstruction of two viral genomes distant from as less as a hundred nucleotides (out of 4450nt length genome, i.e. 2-3%), and present within artificial mixes at up to a 99/1 ratio. Then, we used this method to subsequently analyze mixed infections from field samples, revealing up to three RYMV isolates within one single rice plant sample from Burkina Faso. In most cases, the complete genome sequences were obtained, which is particularly important to better estimate the viral diversity and permits to detect recombination events.

The described methodology consequently allows to identify various haplotypes of RYMV simultaneously infecting a single rice plant, obtain their full-length sequences, as well as a rough estimate of relative frequencies within the sample. It is efficient, cost-effective, as well as portable, so that it could further be implemented where RYMV is endemic. Perspectives include to decipher mixed infections involving other RNA viruses threatening crop production worldwide.

## Introduction

The rise of high-throughput sequencing technologies revolutionizes our vision of global viral diversity, with an exponential increase in the rate of virus discovery in the last decade [1]. Embracing a virome approach (i.e. metagenomics analysis of viruses) led to the characterization of viral diversity in natural ecosystems [2, 3], or to the discovery of zoonotic agents with strong application for human health [4, 5]. For plant viruses, virome approach was applied to both crops and wild plants [6, 7], with different methodologies [8], for example the VANA _Virion-associated nucleic acid-based _ metagenomics [9]. Recently, the methods based on Oxford Nanopore Technology (ONT) gain more attention for viral genomics and metagenomics, thanks to the reduction of ONT sequencing errors, associated with their portability and cheap capital costs [10]. The use of ONT in plant virology is consequently slowly spreading the community of plant virologists, with important applications for virus detection and surveillance, as well as whole-genome sequencing [11–16]; as highlighted in the reviews on ONT and plant virus detection [17, 18]. These innovations have strong implications for risk evaluation of new crop virus emergence and on the rapid development of efficient disease control strategies.

In the case of strong local epidemics with active transmission and co-circulation of various phylogenetic lineages, the probability of a single plant to be simultaneously infected by various viral haplotypes may be high (see for example [19]). Such mixed infections contrast with the case of any (single) viral infection, where within-host replication leads to the formation of viral clouds composed of closely related genetic variants, known as quasispecies [20]. Mixed infections involve more distantly related haplotypes, and in this case, within-plant dynamics of various haplotypes may differ from between-plant dynamics (transmission), with strong consequences for viral evolution [21–24]. Further research on these evolutionary and epidemiological consequences of mixed infections require deciphering the genomes of the distinct viral isolates simultaneously infecting a plant [21, 25]. To this purpose, classical molecular biology techniques (cloning and Sanger sequencing) may be used but are time consuming, while ONT long reads constitute an opportunity to get access to the multiplicity of genotypes within one plant sample and to reconstruct within-plant diversity. However, this issue is not straightforward in terms of bioinformatics analyses, as it requires teasing apart highly similar sequences obtained in the case of distinct genotypes from the same viral species. Recent rise in interest in within-host viral diversity led to the development of various bioinformatics methods on this topic [26–29].

Rice yellow mottle virus (RYMV, genus *Sobemovirus*), causes a rice disease endemic to Africa, that constitutes a threat to the sustainable development of rice cultivation in the continent [30]. RYMV is a (+) single strand RNA virus, with a genome of ca. 4450nt, lacking 3’ polyA tail and organized in five overlapping open reading frames [30]. A recent study evidences a hotspot of rice yellow mottle disease in Burkina Faso [31, 32]. In this irrigated site, a particularly high RYMV genetic diversity was found, with at least four distinct genetic groups co-existing over years [32]. A fraction of samples (6/138, 4.3% of the dataset) analyzed in this study presented mixed Sanger chromatogram (ORF4 gene coding coat protein), suggesting multiple independent infections, resulting in the simultaneous presence of various RYMV genomes within plant.

This study aims at identifying a strategy for ONT sequencing library preparation and data analysis of obtained reads to reconstructing the complete genomes of several distinct viral haplotypes co-infecting a plant. We first performed preliminary tests to assess the best methodology to prepare the libraries for ONT sequencing. Second, we generated artificial mixes to assess the performance of the proposed sequencing and bioinformatic approach. Finally, we applied the validated approach to field-collected samples to decipher mixed infections in the agrosystem. The methodology was also used to obtain full length genomes of greenhouse isolates in a cost and time effective way.

## Methods

### Initial sequencing of single isolate to compare three different ONT sequencing strategies

Three different methods for preparing the libraries were tested. Firstly, the cDNA-PCR Sequencing Kit (SQK-PCS109) from ONT was used. To this purpose, we first performed the reverse transcription (RT), respectively with VN Primer from ONT Kit 5’-5phos/ACTTGCCTGTCGCTCTATCTTCTTTTTTTTTTTTTTTTTTTTVN 3’ and simultaneously with two RYMV specific primers (position 1756 5’ACTTGCCTGTCGCTCTATCTTCCTCCCCCACCCATCCCGAGAATT 3’ and 4446 nucleotides 5’ ACTTGCCTGTCGCTCTATCTTCGGCCGGACTTACGACGTTCC 3’). After the denaturation step, Strand-Switching Primers (SSP from ONT) 5’ TTTCTGTTGGTGCTGATATTGCTmGmGmG 3’ have been added and incubate 2 min at 42°C, followed by 1h at 42°C after the addition of 1 µl of Maxima H Minus Reverse Transcription. The PCR has been done with cDNA Primer (cPRM from ONT) which attach the adapter primer link. We used Rapid Adapter (RAP) to the amplified cDNA library for sequencing with a Flongle Flow Cell R9.4.1 in a MinION MK1C. The sequencing duration ranged from 24 to 48 hours.

Secondly, we performed a direct cDNA sequencing method with the ONT Direct cDNA Sequencing Kit (SQK-DCS109) as described by [16]. Thus, the extracted and DNase treated RNA has been ribodepleted with the QIAseq Fast Select rRNA Plant Kit (Qiagen Hilden, Germany). Double stranded cDNAs were synthesized with random hexamer and the Maxima H Minus Kit (Thermo scientific, K2561). The cDNA products were repaired and dA-tailed followed by ligation with the adapter MIX (AMX from ONT). A Flongle Flow Cell R9.4.1 was used for the sequencing and the library was loaded as recommended by ONT (Flongle Sequencing Expansion Kit EXP-FSE002).

These different approaches were carried out using RNA extracted with the GeneJET Plant RNA Purification Mini Kit (Thermo Scientific, K0802) from rice leaves inoculated by RYMV isolate BF710 collected in the irrigated perimeter of Banzon (Burkina Faso) in 2016 (S1 Table [32]). Each of the three methodologies (VN Primer, RYMV specific primers and direct cDNA) were performed twice, using RNA extracted from plant inoculated with the BF710 isolate.

The basecalling was performed with guppy 6.3.7 [33]. In order to visualize the coverage of RYMV for each run, we use the mapping tool Minimap2 v2.24 [34] and the samtools suite v1.10 [35] with the isolate used for the run as a reference.

### Sequencing artificial mixed-infection samples

Total RNA was extracted using the RNeasy Plant Mini Kit (Qiagen, Hilden, Germany) from rice leaves infected by different RYMV genotypes. To prepare artificial mixes, we chose a set of three RYMV genotypes from Burkina Faso (S1 Table): BF710, BF711 (Banzon, collected in 2016), [32] and BF706 (Karankasso Sambla, collected in 2014) [36]. Full-length genomes of these three isolates were obtained by Sanger sequencing. Their divergence ranges from 2% (110 nucleotides) (BF710 and BF711) to 3% (131 nucleotides) (BF710 and BF706).

A set of five artificial mixed samples was performed, involving two of the three genotypes in different relative proportions (Table 1). To this purpose, estimation of RYMV viral load in samples is a mandatory step, and was done by qRT-PCR according to the protocol described previously with slight modifications [37]. From 1 µg total RNA extract from leaves infected respectively by BF706, BF710, BF711 genotypes, three reverse transcription were performed with the reverse primer R0 (nucleotides 1748–1767: 5′-GGCCGGACTTACGACGTTCC-3’). 5 µl of each cDNA (diluted at 1:1250) were followed by three qPCR (nine qPCR reactions in total per sample) mixed with 12.5 µl of Full Velocity Master Mix (Stratagene) and 300 nM sense and antisense primers (FqPCR2 sense primer, nucleotides 673-690 5’ -ACCTCCTCATCGTCTTGG-3’ and RqPCR2 antisense primer nucleotides 1049-1064 5’ -CGGCGGCACATCTTCG-3’). The number of copies per 1 µg RNA was calculated. Dilutions were performed to adjust the concentrations to 4,3 × 10^−7^ copies. The artificial mixes were then obtained with the desired relative proportions: 50/50 in some cases and 99/1 in others. They were sequenced through the “direct cDNA” methodology, as well as the two initial isolates BF711 and BF706.

**Table 1:**
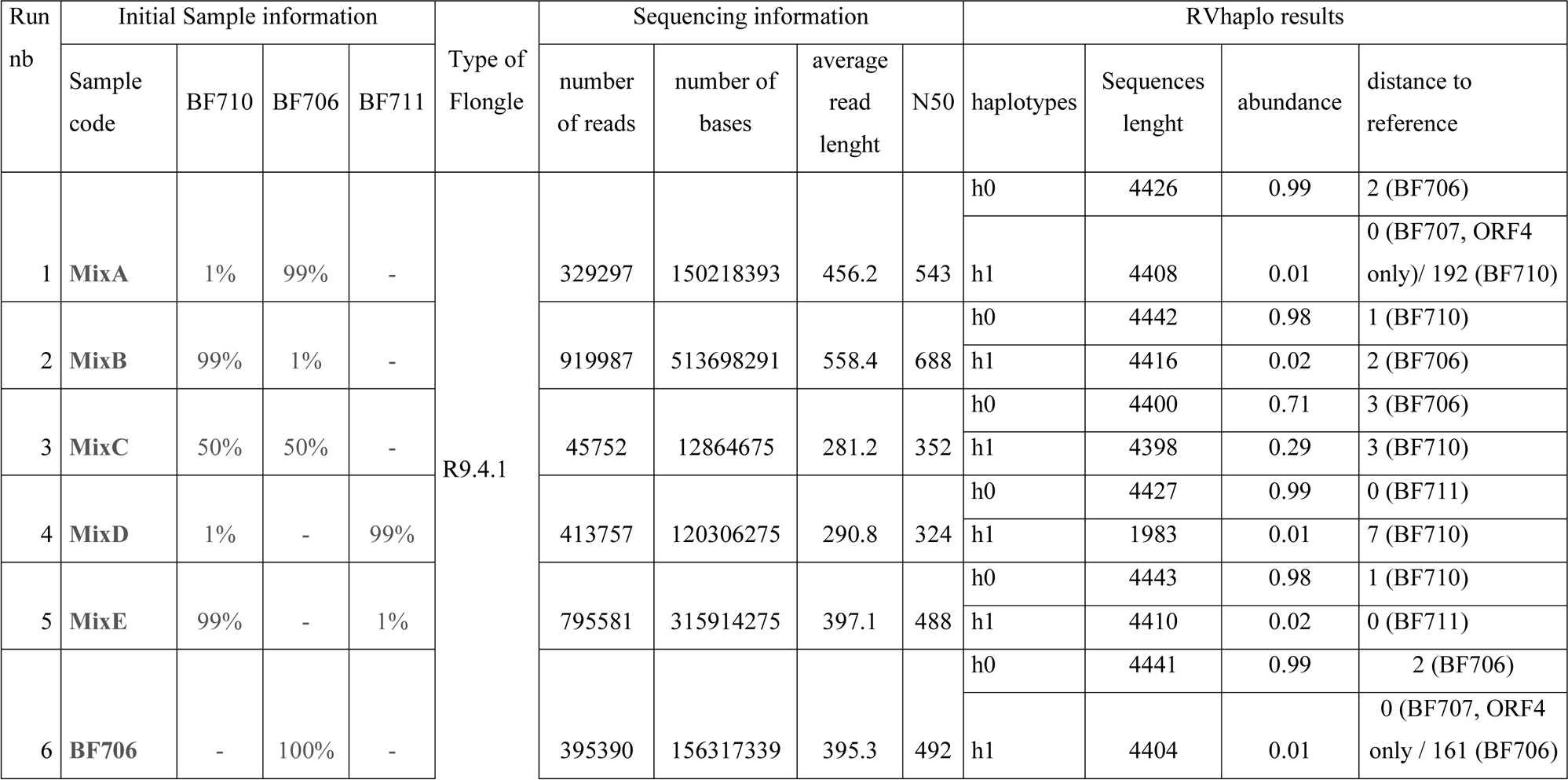
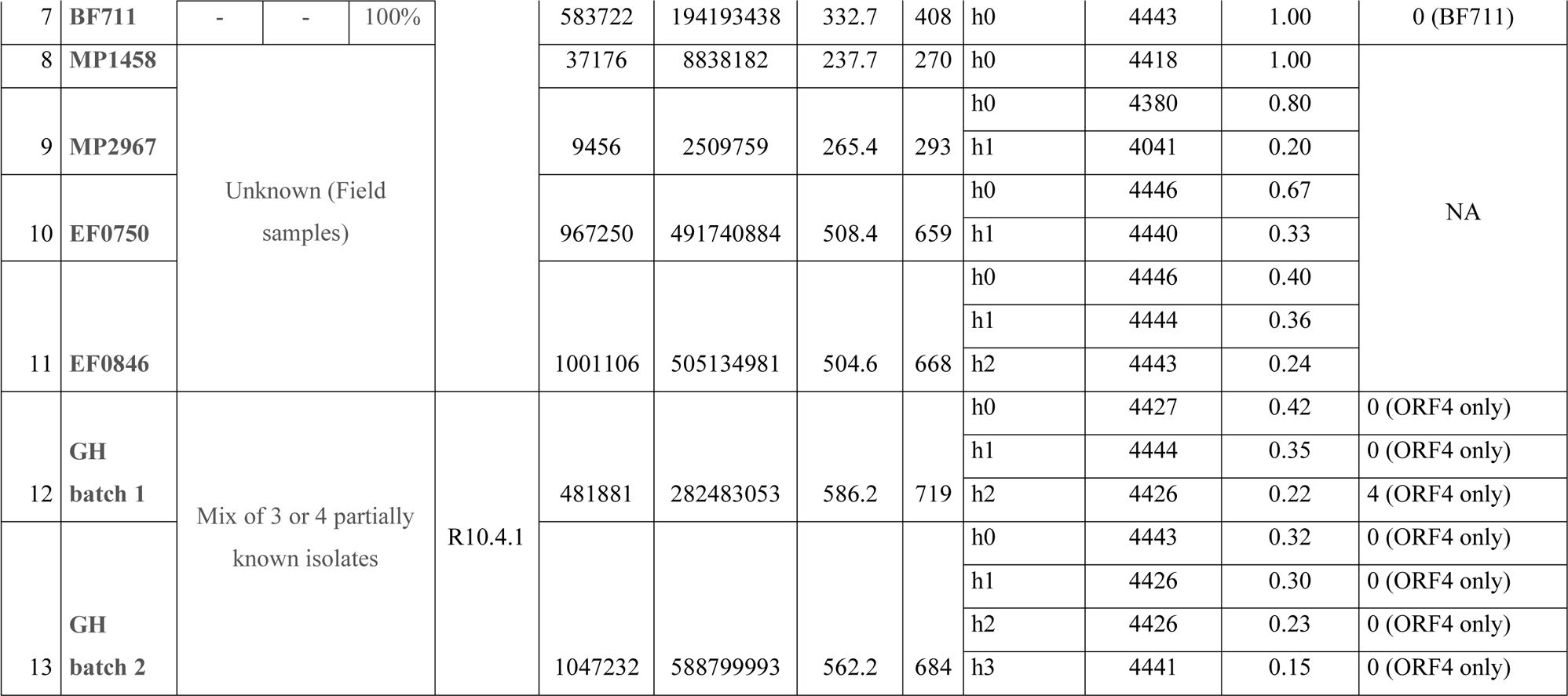
List of the 15 nanopore runs performed by direct cDNA sequencing from RT with random hexamer primers after ribodepletion, and obtained results, in terms of basic sequencing information (after basecalling), as well as haplotype reconstruction (using RVhaplo analysis).

### Haplotype reconstruction from artificial mixes

The basecalling was performed with guppy 6.3.7 [33]. Then, after preliminary assessment of various methods, AssociVar [27], Nano-Q [29], Variabel [38] (data not shown), we chose the bioinformatic tool RVHaplo [26] for reconstructing viral haplotypes from ONT sequencing of mixed-infections.

A snakemake pipeline was made in order to launch several datasets simultaneously (https://forge.ird.fr/phim/acomte/rvhaplosmk) and to favor reproducibility. For each dataset, after initial tests of various thresholds, we chose to remove from the datasets (prior to the analysis) all reads shorter than the N50 (i.e. the length of the read found at the middle point of the length-order concatenation of all obtained reads; or in other words: the length of the shortest read in the group of longest sequences that together represent half of the nucleotides in the set of sequences). This led to speed up the calculations, and obtained data were more consistent with expectations.

### Sequencing field-collected samples

Four rice leaves samples, symptomatic for RYMV, were collected in Banzon irrigated perimeter (south western Burkina Faso) and were chosen for application of the proposed methodology because the amplification of the RYMV ORF4 (coat protein gene) Sanger sequencing presented mixed chromatogram. First, the two samples MP1458 and MP2967, collected in 2017-2018 in Banzon (see doi.org/10.23708/8FDWIE [31] and S1 Table), and already studied in terms of OFR4 sequencing in [32] were chosen. These samples were kept dried right after sampling, and then maintained at ambient temperature [32].

Then, we analyzed two rice leaves samples, EF0750 and EF0846, collected in 2021 (S1 Table), that also presented mixed chromatograms when sequencing ORF4 with Sanger method (unpublished data). Sample conservation was better for these two samples collected in 2021 (they were kept dried and then frozen at −20 a few weeks after sampling), compared to older samples.

### Sequencing greenhouse samples

We then aimed at obtaining full-length genomes of seven RYMV isolates originally collected in Burkina Faso in 2021, and used in greenhouses experiments, for which a partial sequence (ORF4, Coat Protein gene) was available by Sanger sequencing. To this purpose, and to be cost and time effective, we arranged the seven isolates into two batches: EF0252, EF0580, and EF0768 in “Greenhouse batch 1”, and EF0316, EF0321, EF0562, EF0644 in “Greenhouse batch 2”. These isolates all originate from a sampling performed in 2021 in two sites from western Burkina Faso: Banzon and Badala (S1 Table). They were amplified on rice (cultivar IR64) grown in greenhouses.

The Direct cDNA Sequencing Kit (SQK-DCS109) is no longer available since June 2023. As recommended by ONT, we then used their latest Q20+ chemistry and the V14 Ligation Sequencing Kit (SQK-LSK114). We produced double strand cDNAs followed by the adapter ligation using the ONT protocol ligation sequencing kit V14 and used a Flongle Flow Cell R10.4.1 for the sequencing step. We then analyzed obtained data as described previously.

### Phylogenetic analysis of obtained genomes

The different nanopore runs performed led to a total of 28 sequences, all except one being longer than 4 000nt (see Table 1, and S1Fig D). All the 27 almost full-length sequences were aligned with the three reference genomes obtained by Sanger sequencing. Then, a Neighbor-Joining (NJ) phylogenetic tree was built with MEGAX [39] to represent the diversity obtained from analyzed samples.

Such obtained sequences were then attributed to 19 distinct genome sequences, that were then compared with 47 full-length genome sequences from isolates collected in West and Central Africa, which were available in NCBI (July 2023). Maximum-Likelihood (ML) phylogenetic tree were reconstructed using the best-fitting substitution model (GTR+G+I) determined with MEGAX, with 100 bootstrap replicates. Phylogenetic trees were drawn using FigTree v1.3.1 (http://tree.bio.ed.ac.uk/software/figtree/). An estimate of genetic diversity was obtained from MEGA for the newly obtained sequences, compared to the sequences available in the literature.

Finally, potential recombination signals from complete RYMV genome sequences were searched using the seven algorithms implemented in the RDP program v4.101 [40]. Only recombination events detected by at least four methods and with *P*-values below 10^−2^ were considered.

## Results

### Comparing three ONT sequencing strategies

The Fig1 presents obtained results for the three sequencing strategies, in terms of genome coverage, number of reads, read length, proportion of reads mapped on RYMV genome, for the six runs performed on the same RYMV isolate (BF710). The direct cDNA sequencing approach, following [16], yields the highest percentage of reads mapped over the RYMV genome (95% compared to 19-47%, on average 33% for the two other methods). Average read length was also much higher for this method (on average 648 nucleotides) compared to others (335-423 nucleotides). Although the total number of reads obtained was lower than when using VN Primers (oligo-dT) technique, the direct cDNA sequencing gave by far the best results in terms of genome coverage (Fig 1) and was chosen for the following runs.

**Fig 1:**
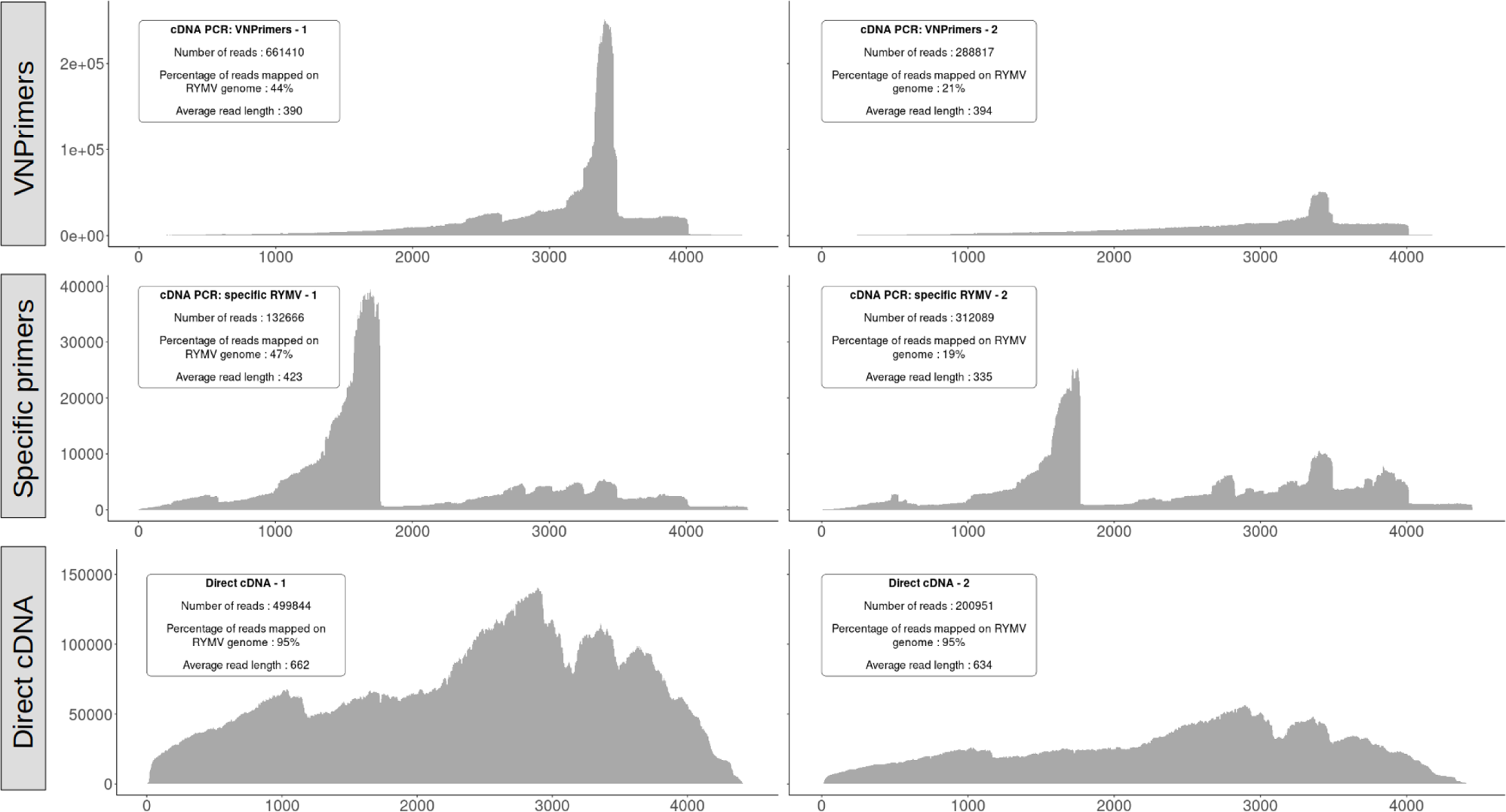
Comparison between the three methodologies for RYMV sequencing of the BF54 isolate, with two Oxford Nanopore Technology (ONT) sequencing replicates (left and right panels of the figure) for each of the three methods: amplification with VN Primers on top, RYMV specific primers on the middle, and direct CDNA sequencing on the bottom.

It is to note that RYMV often encapsulates also a small circular satellite RNA [30], likely present in our samples as it was reported to be present in all isolates originated from West and Central African [41], but this 220nt long sequence was not found in our ONT sequences (data not shown).

### Analyzing artificial mixes results to assess the bioinformatic pipeline

We then tested whether we could find back the different RYMV haplotypes in artificial mixes of two RYMV isolates. To this purpose, we used two additional RYMV isolates (BF706 and BF711), also originating from Burkina Faso, in addition to the reference isolate BF710 mentioned before. The Table 1 presents obtained results for the five artificial mixes and the two isolates BF706 and BF711, all performed with the direct cDNA sequencing approach.

We sequenced samples (mixA-mixE) obtained from these isolates, previously quantified, and mixed in different relative proportions of two isolates: equal proportion (50-50%, mixC), or a major isolate in mixed infection (99-1% mix A, B, D, E) (these artificial mixes correspond to runs 1-5 in Table 1). We then sequenced the two isolates BF711 and BF706 taken separately (see runs 6-7 in Table 1). For these 7 runs, the number of reads ranged from 45 752 to 919 987 (mean = 497 640, Table 1 and S1 Fig B), and the average read length from 281 to 558 (mean = 387) (Table 1).

For the artificial mixes (runs 1 to 5), RVHaplo managed in most cases to identify the multiplicity of infection and to reconstruct the genomes. For the 50/50 (mixC, run 3), we obtained the two expected genomes (ca 4400nt) with only 3 differences from Sanger sequences in each, and RVhaplo estimated the relative proportions to 70/30 (Table 1). For the other mixes, all with relative proportions of 99/1, we obtained in most cases the expected haplotypes, with full length genome reconstruction and relative estimate 99/1 or 98/2. This is the case for mixB (run 2) and mixE (run 5). For mixD (run 4), it was only partially true since the minority haplotype was not complete, nor perfectly matching (BF710: 1983nt, 7 errors); a result most likely attributable to the lower quality of this specific run (N50 = 324, Table 1 and S1 Fig C).

On the other hand, for mixA, we obtained the sequenced of the majority isolate (BF706), but the minority (1%) haplotype does not correspond to the minority haplotype put in the mix (BF710). Instead, its ORF4 partial sequence corresponds exactly to another isolate manipulated in greenhouses at the same time (BF707; [36]). When sequencing the isolates independently (runs 6-7), we obtained exactly the expected haplotype (4443nt) for the isolate BF711 (run 7), but two distinct isolates for BF706 (run 6). Indeed, the first haplotype corresponds to the expected isolate BF706 (with only 2nt difference out of 4441nt total), while the second unexpectedly corresponds a haplotype distant from 161nt from BF706 (run 6), but matching the BF707 isolate over the ORF4 partial sequence. We consequently interpret the results obtained for both run 1 and 6 as a contamination of BF706 by BF707 occurred during leaf manipulations (as these two isolates were not manipulated together during the lab stage, but in the greenhouse instead). For MixA (run 1), the minor isolate (BF710) could be retrieved when changing the threshold of read length applied prior to data analysis (set to the N50 for all the analyses presented here). Indeed, when the threshold was set up 700 instead of 543, three haplotypes were deciphered: BF706 (98%), BF707 (1%) and BF710 (1%), all three with minimum length 4336, and maximum 2nt difference from Sanger reference.

Overall, we appreciate the efficiency of approach in deciphering the haplotypes present in artificial mixes, so that these results validate the approach for library preparation and bioinformatics analysis. The phylogenetic tree (S1 Fig E) clearly shows a difference between the different haplotypes (distant from 110 to 192nt, 2-4% of the genome), compared to artefactual haplotypes (maximum 7nt difference). These runs of known composition consequently allow to estimate a range of threshold for the divergence of haplotypes that could be deciphered through the approach: mixed infections involving genomes distinct from 2% should be retrieved by the proposed methodology, while it would most likely not work out for genomes differing for less than 0.05% of their genomes.

### Deciphered mixed infections and newly obtained full length genomes

Once the sequencing strategy and the bioinformatic tools were validated, we applied the approach to a set of four field samples, i.e. rice leaves collected in Burkina Faso. While the two samples collected in 2017-2018 (MP1458 and MP2967, runs 8-9 Table 1) gave lower quality results (average read length maximum 265), most likely as a consequence of a conservation issue, each of the two samples collected in 2021 (EF0750 and EF0846, runs 10-11) led to good overall sequencing (ca 10^6^ reads, N50 approx 650nt, see Table 1, S1 Fig B-C). While one sample (MP1458) only led to one haplotype, the three others led to various haplotypes (differing from 110 to 265nt within each sample), with up to three complete (minimum 4443nt) genomes in proportions 40/36/24 retrieved from the sample EF0846 (run 11, Table 1). Compiling the results of these four runs performed on field samples, we obtained a total of eight new RYMV genomes that could be placed in a phylogenetic tree (Fig 2, six haplotypes in green). It is to note that one of these haplotypes (2018MP2967_h1) was shorten in its 5’ part as a consequence of a lower quality area (10 gaps in ca 150nt area) detected when aligning to the sequences from the literature. This haplotype is the minority one from a lower quality sample and all other sequences were fine.

**Fig 2:**
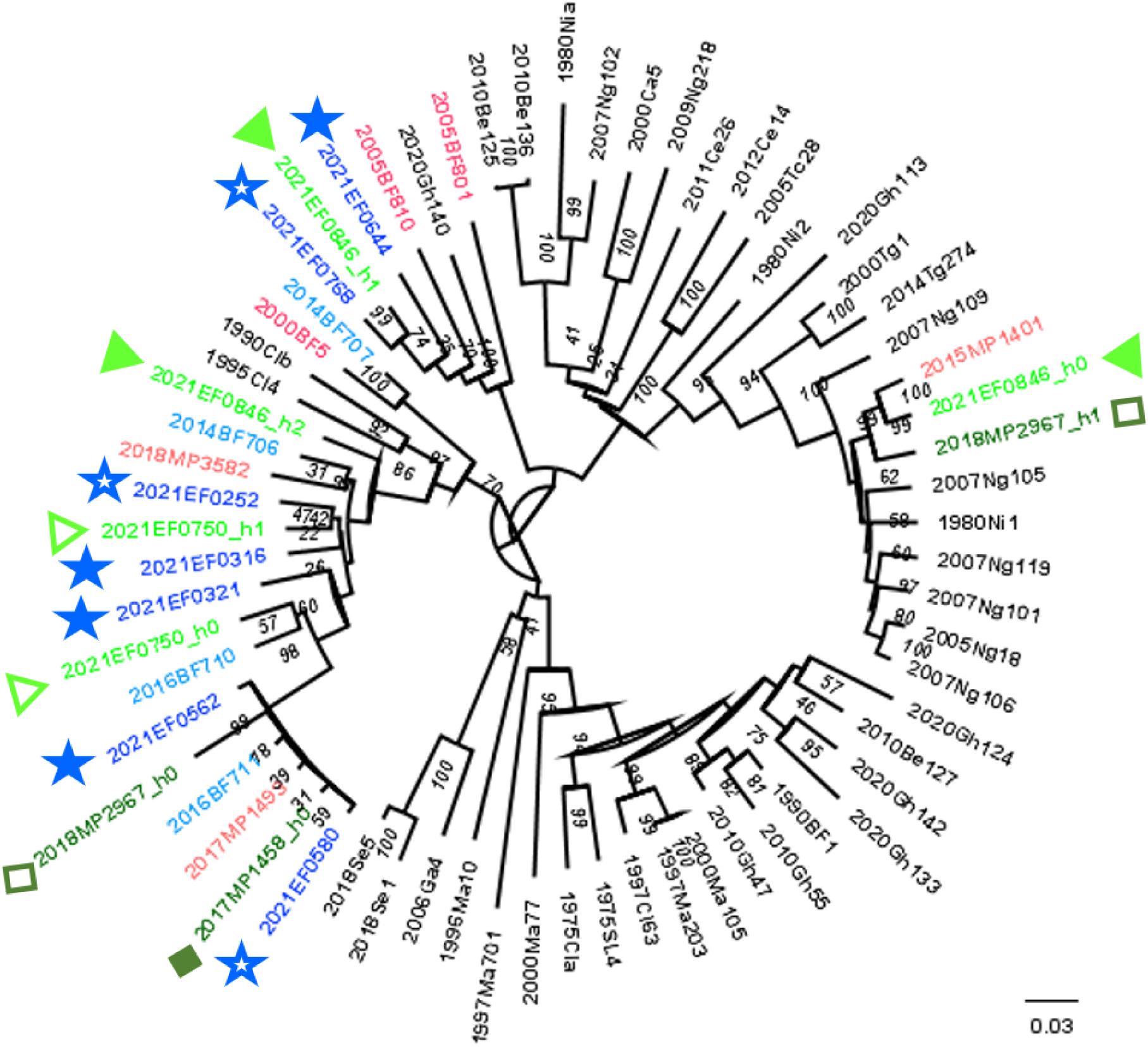
Phylogenetic tree replacing the 19 RYMV haplotypes obtained by nanopore sequencing (this study) within the 47 viral genomes already published from samples collected in West Africa (literature). Maximum-Likehood (ML) phylogenetic tree of 47 RYMV genomes of isolates from West- and West-Central Africa (NCBI database) along with 19 haplotypes from this study. The ML phylogenetic tree was built based on the full-length genome sequences (4461nt). The isolates from greenhouses are colored in blue (light blue for the isolates sequenced as single, and dark blue for the batches), while the field isolates are represented in green (dark green for samples collected in 2017-2018, and light green for others, collected in 2021). Each symbol corresponds to a single nanopore run, indicating the different haplotypes obtained for that particular run, except for the isolates used as references (BF710, BF711, BF706, as well as the unexpected BF707). The samples from the literature also originating from Burkina Faso are labelled in orange.

In addition, we used the same strategy to obtain seven full-length genomes from two Flongle runs (runs 12 and 13, Table 1) performed on artificial batches comprising three to four isolates. Obtained sequences were analyzed using the developed pipeline, resulting in the expected number of distinct haplotypes for both runs. By comparing obtained genomes to partial ORF4 sequences (Sanger sequencing), we could retrieve full-length genomes of the seven distinct isolates (Fig 2, haplotypes in blue, labeled with stars).

The phylogenetic analysis of 66 genomes (Fig 2) shows the contribution of these nanopore sequences to the RYMV genomic diversity in West Africa: 19/66 = 28.8% in terms of number of viral genomes. Although the sampling is local (southwestern Burkina Faso), this area being a hotspot of RYMV diversity [32], this dataset constitutes a substantial contribution to the genetic diversity of RYMV in West Africa. Indeed, the genetic diversity of these 19 new sequences is 0.043 ± 0.002 substitution/site, compared to 0.065 ± 0.003 subst./site for the 47 sequences from the literature (overall genetic diversity: 0.062 ± 0.003 subst./site). Finally, the recombination analysis (RDP4, see S2 Table) detected one recombination event newly evidenced in Burkina Faso, in addition to the S1bzn group notified in [32], so that at least two recombinant haplotypes circulate within the irrigated perimeter of Banzon.

## Discussion

The local co-circulation of multiple phylogenetic lineages, and within-plant viral diversity is particularly likely for rapidly evolving pathogens such as RNA viruses [42]. Mixed infections, and subsequently virus-virus interactions, are likely to affect viral diversity and evolution [43]. In such context, long reads generated by ONT appear as promising to decipher the multiple viral genomes simultaneously infecting the same host. Indeed, our results show that it is possible to use direct cDNA ONT sequencing combined with the RVhaplo bioinformatics tool, to identify various haplotypes of the rice yellow mottle virus (RYMV) co-infecting a single rice sample, and to obtain their full-length sequence. This was true even for 99/1 mixes of isolates, that differ only from 2-3% of their genomes (approximately a hundred nucleotides out of a ca 4450nt viral genome).

The methodology described in Liefting et al [16], based on cDNA direct sequencing, efficiently yielded almost full-length RYMV genome, with obtained sequences either exactly, or very closed (i.e. 2nt out of 4450nt genome) to the same samples sequenced with Sanger method. The proportion of reads mapping on RYMV genome was much higher with this method (95%) compared to PCR-based strategies, both using VN primers, or RYMV specific primer (range from 19 to 47% of reads mapped on viral genome). Average read lengths were also higher with the cDNA direct sequencing strategy, that was consequently selected to go further in sequencing mixed-infected samples.

We note however that the sequencing method based on reverse transcription with polyT primers works out even though the RYMV has no 3’ polyA tail. This was also observed for other viruses (as for Potato leafroll virus [14]). Here, this result can be explained as the repartition of reads along the RYMV genome is highly aggregated on A-rich regions of the genome (data not shown). Although this strategy was left up for the purpose of this study, the information on the possibility to sequence RYMV from polyT primers RT may be useful, e.g. for metagenomics studies.

The RVhaplo bioinformatic tool [26] was used to analyze the data obtained through cDNA direct sequencing (see above), and allowed to efficiently reconstruct the different haplotypes present as mixed infections in our study. This was true even in the case of mixed infection involving a mix of a majority isolate at 99%, mixed with only 1% of another isolate, demonstrating the sensibility of this approach. Also illustrating the high sensibility of the methodology, is the detection of an unexpected haplotype in one of the isolates multiplied in controlled conditions (greenhouses). Only representing 1% of the analyzed sample, this haplotype would likely remain undetected using Sanger sequencing, so that an application of the methodology presented here is also to screen a sample for potential contamination prior to setting up sensible greenhouse experimentations.

The described methodology is quick and appears as cost-effective. Indeed, in this work, the use of two Flongles led to seven genomes for greenhouse isolates, and we also generated five genomes from two Flongles used on two field samples (similar pricing rate compared to Sanger in our conditions). Globally, this study generated 19 full-length genomes, compared to 47 available to date in total for RYMV in West Africa, so that it expands the knowledge in terms of full-length genomes in this area. This is particularly relevant in the context of viral diversity hotspots, as is the case for western Burkina Faso, where distant genetic groups are co-circulating, giving the chance for recombination [32]. Other recombinant isolates were previously reported for RYMV in East Africa [44], the center of origin of RYMV, where RYMV genetic diversity is highest. Also, viral epidemics in Madagascar were shown to result from a unique introduction of a lineage potentially resulting from a recombination event [45]. Recently, increased RYMV dispersion led to lessen the historically strong geographic structuration of viral diversity in West Africa [32, 46, 47]. Such a context is likely to enhance the risk of recombination between distant genetic lineages. In western Burkina Faso, a recombinant lineage was already evidenced in a previous study [32], and the additional genomes obtained in this study allowed to identify a second recombination event. Further ONT sequencing of RYMV genomes likely would lead to detect others, especially as the period of co-circulation of lineages gets longer. This potentially increases risks in terms of resistance breakdown and pathogenicity [48].

Deciphering the different genomes simultaneously present within a host in the context of mixed infection remains a methodological challenge. Our study managed to decipher mixed infections of rice infected by various RYMV isolates, even for mixes in 99/1 proportion, and two isolates that only differ from 2-3% of their genomes (ca one hundred nucleotides out of a ca 4450nt viral genome). This was most likely allowed by the use of a long-read sequencing approach, as also reported with the detection of two strains of DNA virus [11], or using PacBio long-read HiFi sequencing to unravel intra-host diversity across natural RNA mycovirus infections [49]. Previous attempts to characterize within plant diversity in RYMV using a short-read strategy (Illumina technology) allowed to identify that the sample corresponds to a mixed infection, and estimate the proportion for each polymorphic site, but it could not phase the polymorphic sites and reconstruct the distinct viral genomes (Poulicard et al, unpublished). On the other hand, sequencing virus-derived small interfering RNA (vsiRNA) led to distinguish several strains from three distinct viral species in strawberry [50]. A recent systematic comparison of ONT and Illumina sequencing from total RNA (unfortunately not including the cDNA-direct sequencing method applied in this study) showed the performance of ONT sequencing in terms of quality, affordability and practicality, for different viral organization and including low titer virus [51]. In this study, the high titer of RYMV in rice (strong within plant accumulation [37]) may have helped, and the applicability of the described methodology to other RNA viruses remains to be evaluated.

Such mixed viral infections are important as interactions between viral genomes (‘social life of viruses’, [24]) may affect plant disease ecology and evolution [21, 25]. For fungi, Lopez-Villavicencio et al [52] showed that fungal genotypes found in each plant are more related than expected by chance, suggesting that *Microbotryum violaceum* actively excludes dissimilar genotypes while tolerating closely related competitors. The methodology described in this article allows to distinguish genomes distant from 110nt out of ca 4450nt (2%). On the other hand, it does not capture the viral clouds of highly related genetic variants generated by within-host replication in high mutation rate RNA genomes. Instead, it allows to focus on supposedly independent infections by various RYMV isolates.

ONT sequencing is a highly dynamic methodology (as exemplified here with the change from flow cells R9 to R10 during this study). Its flexibility and diversity of library preparation make it capable to adapt to different context (as a ‘swiss knife’). This combined to its practicality and performance renders ONT sequencing an ideal tool for fast and affordable virus detection and genomics [51]. It will continue to evolve, and with no doubt improve in the next future, making it a highly promising technology not only for viral genomics but in general [53]. As a portable sequencing technology [12], not requiring costly initial investment and cost-effective compared to Sanger or Illumina, it holds promise for the global South, and particularly in Africa. Further deployment of ONT sequencing in Africa requires capacity building, mostly in terms of bioinformatic resources and skills, and various initiatives are under way to achieve this goal (in Mali [54], or Burkina Faso, pers.com.). Such capacity building is required to further understand and fight crop diseases all around the world, a matter of the utmost importance considering that highest losses frequently corresponds to emerging diseases in food-deficit regions with fast-growing populations [55].

## Acknowledgments

This work was performed within the formalized partnership “International joint Laboratory LMI PathoBios: Observatory of plant pathogens in West Africa: biodiversity and biosafety” (www.pathobios.com; twitter.com/PathoBios). We thank the rice farmers from the study sites in Burkina Faso for their kind collaboration. We also thanks Dr Cai for helpful advices at early stages of bioinformatic analyses. The manuscript benefited from helpful comments from Eugénie Hébrard. The authors acknowledge the ISO 9001 certified IRD itrop HPC (member of the South Green Platform) at IRD Montpellier for providing HPC resources that have contributed to the research results reported within this paper, URL: https://bioinfo.ird.fr/, http://www.southgreen.fr

This work was publicly funded through ANR (the French National Research Agency) under the project EVCOPAR (ANR-20-CE35-0004-01).

## Supporting information

**S1 Table.** List of RYMV-infected rice leaves samples from Burkina Faso analyzed in this study: site and field of origin, GPS coordinates and date of sampling.

| Code | Site | Field ID | GPS coordinates | Sampling date | Analyzed as |
| --- | --- | --- | --- | --- | --- |
| BF710 | Banzon | na | 11.321 ; -4.805 | 17/10/2016 | ‘greenhouse isolate’ |
| BF711 | Banzon | na | 11.322 ; -4.805 | 17/10/2016 |  |
| BF706 | Karankasso<br>Sambla | na | 11.247 ; -4.563 | 07/10/2014 |  |
| MP1458 | Banzon | BZ11 | 11.335 ; -4.794 | 12/10/2017 | ‘field sample’ |
| MP2967 | Banzon | BZ09 | 11.336; -4.790 | 15/11/2018 |  |
| EF0750 | Banzon | BZ21 | 11.323; -4.803 | 08/11/2021 |  |
| EF0846 | Banzon | BZ23 | 11.330; -4.809 | 09/11/2021 |  |
| EF0252 | Badala | BL22 | 11. 368; -4. 373 | 05/10/2021 | ‘greenhouse isolate’ |
| EF0316 | Badala | BL24 | 11.387; -4.363 | 13/10/2021 |  |
| EF0321 | Badala | BL24 | 11.387; -4.363 | 13/10/2021 |  |
| EF0562 | Banzon | BZ04 | 11.328; -4.806 | 08/11/2021 |  |
| EF0580 | Banzon | BZ04 | 11.328; -4.806 | 08/11/2021 |  |
| EF0644 | Banzon | BZ08 | 11.336; -4.802 | 09/11/2021 |  |
| EF0768 | Banzon | BZ21 | 11.323; -4.803 | 08/11/2021 |  |

**S2 Table.**
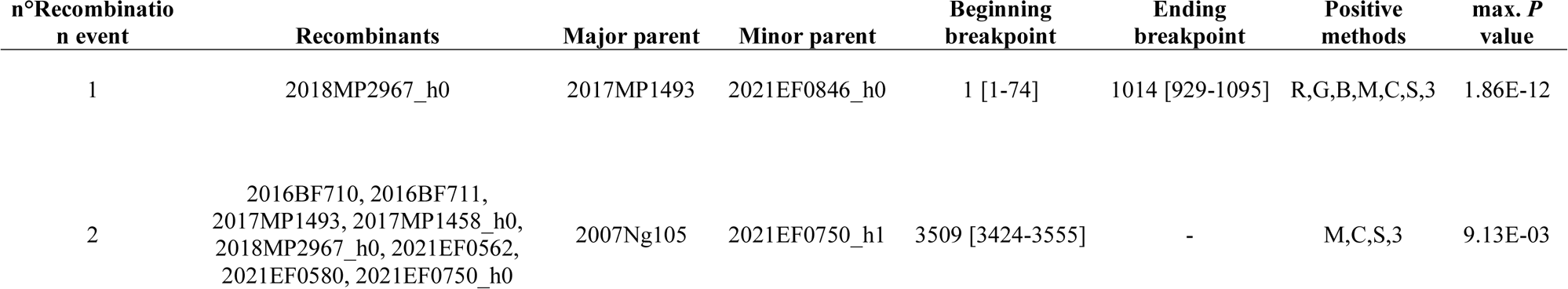
Results of recombination detection using RDP4.10 software. Two recombination events were detected, one in only one RYMV genome, and the other, already documented in Billard et al 2023 [32], is found in eight viral haplotypes. For each of these two recombination events are indicated the most likely parents (major and minor depending on the proportion of genomes attributed to each parental genomes), the breakpoints, the different methods to detect recombination, and the global P-value associated.

**S1 Fig.**
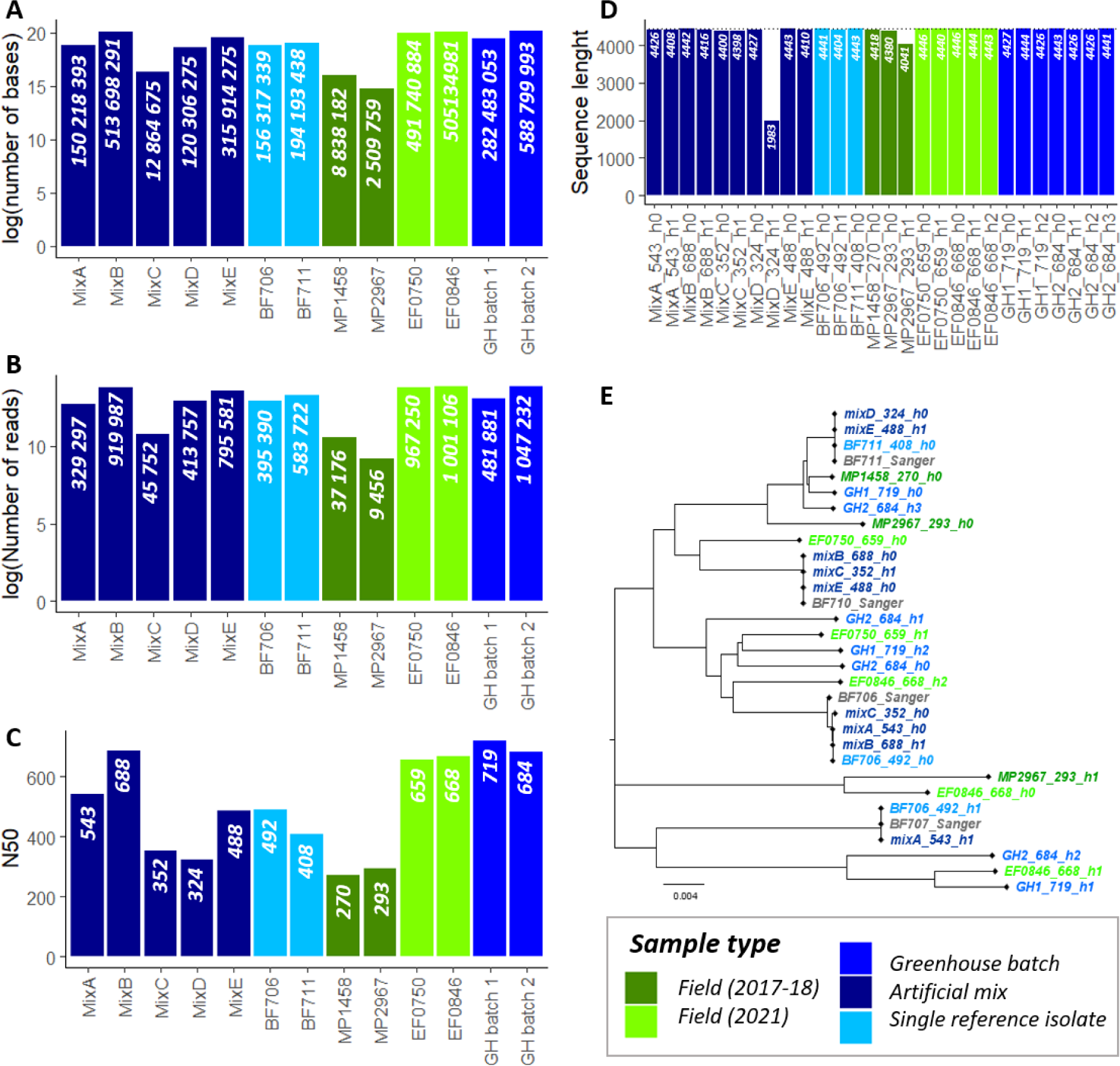
Results of the 13 ONT runs performed though cDNA direct sequencing strategy (see also Table 1). **A.** Total number of bases obtained for each run (log scale). **B.** Total number of reads obtained for each run (log scale). **C.** N50 for each run (defined as the length of the shortest read in the group of longest sequences that together represent half of the nucleotides in the set of sequences). **D.** Sequence length of the 28 different haplotypes reconstructed by RVhaplo method over the 13 runs. **E.** Neighbor-joining phylogenetic tree of the 27 (almost full-length) obtained haplotypes, with the four reference genomes obtained by Sanger technique. The legend, common to the different panels, appears on the bottom right and distinguishes: in green the field samples, and in blue the samples corresponding to viral amplification in greenhouses (and then either sequenced in isolation, or in mixes).

